# ProteinGLUE: A multi-task benchmark suite for self-supervised protein modeling

**DOI:** 10.1101/2021.12.13.472460

**Authors:** Henriette Capel, Robin Weiler, Maurits Dijkstra, Reinier Vleugels, Peter Bloem, K. Anton Feenstra

**Author notes:** these authors contributed equally to this work (Henriette Capel, Robin Weiler, Maurits Dijkstra).

## Abstract

Self-supervised language modeling is a rapidly developing approach for the analysis of protein sequence data. However, work in this area is heterogeneous and diverse, making comparison of models and methods difficult. Moreover, models are often evaluated only on one or two downstream tasks, making it unclear whether the models capture generally useful properties. We introduce the ProteinGLUE benchmark for the evaluation of protein representations: a set of seven tasks for evaluating learned protein representations. We also offer reference code, and we provide two baseline models with hyperparameters specifically trained for these benchmarks.

Pre-training was done on two tasks, masked symbol prediction and next sentence prediction. We show that pre-training yields higher performance on a variety of downstream tasks such as secondary structure and protein interaction interface prediction, compared to no pre-training. However, the larger base model does not outperform the smaller medium. We expect the ProteinGLUE benchmark dataset introduced here, together with the two baseline pre-trained models and their performance evaluations, to be of great value to the field of protein sequence-based property prediction.

**Availability:** code and datasets from https://github.com/ibivu/protein-glue

## Introduction

Machine learning methods have the capability to predict many useful properties of proteins directly from their sequences^1–3^. However, these methods require *labeled* data, mapping proteins to the property of interest. High-quality labeled data is expensive to acquire—and large quantities are usually required in order to train a good predictor.

In the domain of natural language processing (NLP), the issue of missing or scarce labels is often solved by *pre-training* a model on unlabeled, general domain data. This results in representations of the data that capture high-level semantics. These can then be used in *downstream tasks*: specific prediction tasks for which only a limited amount of labeled data is available. Usually, this is achieved by fine-tuning the pre-trained model on the labeled data^4,5^. Training the downstream tasks is therefore also called the fine-tuning step. Recently, the transformer architecture has emerged as a firm favorite for this kind of approach. Transformer models such as BERT^6^ and GPT-2^7^, have shown a remarkable ability to generalize across domains. It is a reasonable question to ask whether this approach carries over to the domain of proteins: can we successfully pre-train a transformer model on unlabeled data, and fine-tune for *a variety* of tasks requiring labeled data. If so, does the pre-training allow us to perform better than if we had trained on the labeled data alone?

Several recent studies have already investigated protein representation models, among which models based on the transformer architecture^8–11^. In general, the representation models make use of a large set of protein sequences to train an NLP based model of which the objective is to learn embeddings that represent the proteins. We provide a comprehensive overview of the protein representation models in the related work section. Most of these models use different protein sequence databases for pre-training, widely differing numbers of model parameters, as well as different downstream tasks for evaluation. Most importantly, these models are generally evaluated only on one or two downstream tasks, giving a poor indication of how well the pre-trained representations generalize. We suggest that the rapid progress of pre-trained transformer models in the domain of NLP is not just due to the power of the models, but also to the wealth of benchmarks for a variety of tasks already available. Without standardized and varied benchmark suites like GLUE^12^, we believe that development would have progressed much slower.

In this paper, we present a benchmark set for the domain of protein prediction, including a variety of structural protein prediction tasks, in order to generalize pre-trained representations, called the Protein General Language (of life) representation Evaluation (ProteinGLUE) benchmark. This benchmark consists of seven downstream tasks, with data formatted for and tested in large transformer models. The following tasks are included:

### Secondary structure

The secondary structure describes the local structure of a protein, defined by patterns of backbone hydrogen bonds in the protein^13^. Commonly, these are three types: *α*-helix and *β*-strand, and anything else is labelled coil; a further subdivision can be made into eight types^14^. Secondary structure prediction methods aim to classify the type per amino acid^2^.

### Solvent accessibility (ASA)

For every amino acid in a protein, the solvent accessibility indicates the amount of surface area of the amino acid that is accessible to the surrounding solvent^15^. Relative solvent accessibility classifies residues into a buried or non-buried state, reducing the difficulty of the prediction task. We model the absolute solvent accessibility as a regression task over every amino acid and the relative solvent accessibility as a classification task over every amino acid.

### Protein-protein interaction (PPI)

The interactions between proteins is arguably the most important property for functioning of a protein^16^. Almost all processes occurring in a cell are in some way dependent on protein-protein interactions; these include DNA replication, protein transport and signal transduction^15,17^. The protein-protein interaction interface determines which residues are involved in the interaction, and may be predicted as a classification task over every amino acid.

### Epitope region

A specific kind of PPI is the binding between antigens and antibodies. The antigen region that is recognised by the antibody is a set of amino acids on the protein surface, and is known as the *epitope*^18^. These may be predicted as a classification task over every amino acid^19^.

### Hydrophobic patch prediction

A number of adjacent hydrophobic residues on the protein surface is called a *hydrophobic patch*. Hydrophobic residues on the surface of the protein may be important for the interaction between proteins, or for membrane interactions, and have been implicated as being a driving factor in protein aggregation^20^. Protein aggregation in turn is thought to be a major causative factor in the development of diseases like Alzheimer and Parkinson.^21^

To estimate performance on these tasks, we pre-trained two large transformer models. This allows us to test our benchmark suite, provide reference code for the training and development of prediction methods, and to give a baseline performance for each task, showing what performance can be expected from a modest sized model. Commonly, pre-training is performed on a large set of unlabeled general domain data. In the protein domain this translates to unlabeled protein sequences. We have chosen to use the protein sequences from the protein domain family database Pfam, which is a widely used database for the classification of protein sequences^22^.

Our pre-training models are based on the BERT transformer architectures for natural language processing^6^. We trained two models of different sizes: the `medium` and `base` model architectures. The `base` model was first described in the paper of Devlin et al.^6^ and contain 12 hidden layers, 12 self-attention heads, a hidden size of 768 and 110M parameters. The smaller `medium` model could be used to overcome the time- and memory limits that are associated with the `base` models^23,24^. This `medium` version contains 8 hidden layers, 8 attention head, a hidden size of 512 and 42M parameters.

Our contributions are as follows:

- A set of generally usable downstream tasks for evaluating pre-trained protein models.
- A repository of reference code showing how to pre-train a large transformer model on unlabeled data from the Pfam database, and how to evaluate it on our benchmarks.
- Two such pre-trained models, broadly similar to the BERT-`medium` and BERT-`base` models.

The rest of the manuscript is organised as follows. We will first give an overview of **related work**, followed by the outline of the **ProteinGLUE Benchmark suite** we present here. We then first describe methods and present results of the **pre-trained models**, and then methods and results of the **fine-tuned models**.

All datasets, code and models are publicly available.^*^ All code is MIT licensed, the models are public domain (that is, creative commons CC-0) and the datasets are each released under the most permissive licence allowed by the source data.

## Related work

Many learning architectures have been used for a variety of protein property prediction tasks. As a background to the multi-task benchmark suite we present here, we therefore first include a rather in-depth overview of relevant natural language modelling learning architecture. We will then proceed to review state-of-the-art machine learning approaches to protein modelling, from which we have gleaned relevant and interesting prediction tasks, sources of reference data as well as some inspiration on our learning approaches. Note that this section provides background information and is not required for the main understanding of this study.

### Natural language modeling

The idea of combining labeled and unlabeled data has a long history in machine learning, going back to such approaches as self-training and co-training^25–27^. One of the first deep learning models to combine unsupervised pre-training with supervised fine-tuning in multiple domains was the RNN-based ULMFit.^28^ This was followed by ELMo^29^, which used bidirectional RNNs and was the first to show state-of the art performance across many downstream tasks. We have recently investigated a number of different Neural Net architectures for their ability to predict protein interfaces^30^. *Transformer* models are architectures which rely primarily on the *self-attention* operation. Self-attention is a sequence-to-sequence operation in which the output vector for each token *i* in the sequence is a weighted average of all input vectors in the sequence, with the weights determined dynamically by the contents of the corresponding input vector. Most commonly, the weight for input *j* is based on the dot product of input vectors *i* and *j*. A key property of transformers is that self-attention is the only operation that mixes information *between* tokens in the sequence. All other operations in the model are applied to each token in isolation.

The Transformer was introduced in by Vaswani et al.^31^. This model was an encoder/decoder architecture designed specifically for machine translation. Devlin et al.^6^ simplified the model to a single stack of transformer blocks, and adopted the pre-training and fine-tuning approach from ULMFit and ELMo. The result, called BERT (Bidirectional Encoder Representations from Transformers), is what we base our reference models on. BERT is a bidirectional sequence-to-sequence model: both its input and its output are sequences of vectors, of the same sequence length. For the computation of each of the output vectors, all vectors of the input may be used (to the left or the right). By contrast, autoregressive models, like those in the GPT family^7,32^ are *unidirectional*, which means that the output vector for one element in the sequence is computed using only the input vectors of preceding elements. Both bidirectional and autoregressive models have been shown to be capable of learning strong representations both of the tokens in the sequence and of the sequence as a whole. In NLP, text sequences are most commonly broken up into tokens larger than individual characters, but smaller than individual words. In the protein setting, individual amino acids are usually taken as tokens.

### Protein modeling

In recent years, attempts have been made to automatically generate enriched embeddings of protein sequences through machine learning. These embeddings generally capture information that is not explicitly encoded in the protein sequence, such as information about the structure or dynamics. This enriched representation can be used in place of the original sequence to improve performance on a variety of tasks, including protein database searching, regression and classification tasks.

Enriched protein representation models can be categorized on three axes. Firstly, there is the architecture of the model used to generate the representation, which can broadly be categorized as either Word2Vec-based^33–35^, LSTM-based^36–40^, or Transformer-based^8–11^.

Although not always specified, a second categorization can be made in terms of the model size, or the number of parameters a model contains. This is to some degree correlated with the model type, but not in an absolute sense. Comparisons between types of models are therefore complicated, if for example more recent Transformer models with many parameters are benchmarked against smaller LSTM models. The LSTM-based representation method UniRep^36^ contains 18 million parameters, the TAPE Transformer-based model contains 38 million parameters^8^, and the LSTM-based model DeepSeqVec contains 93 million parameters^37^. Recently developed methods have mostly been based on Transformer models, and a clear trend can be observed of increasing size. For example, ESM-1b has 650 million parameters^41^, and Prot-TR-XL-UniRef has 3 billion^11^. It is shown that even these large models, such as ProGen with 1.2 billion parameters, still do not overfit the training data^10^. Heizinger et al.^37^ note that the risk of overfitting is generally very small, given the fact that the number of tokens in the training set (hundreds of billions) is much higher than the number of parameters; however for protein data one must take into account the redundancy of very many similar (homologous) sequences. For (large) protein sequence alignments this is commonly done by evaluating the so-called effective number of sequences *N*_*e f f*_ ^42^, essentially counting clusters of sequences at a set threshold of similarity (e.g. 62% or 80%). However, we do not attempt to translate this approach to sequence databases here.

Thirdly, the size of the datasets used for pre-training differs significantly across methods. To give an idea of the range: UMSDProt^38^ is pre-trained on 560k Swiss-prot sequences^43^, UniProt is trained on 24 million UniRef50 sequences, EMS-1b is pre-trained on 250 million UniParc sequences, and Elnaggar et al.^11^ even train their models on the BFD dataset containing over 2 billion sequences^44,45^, and on the UniRef100 database with 216 million sequences. We find that UniRef50^43^ and Pfam are the most popular pre-training databases. By selecting protein (domain) sequences from each cluster or family in Pfam, it can be ensured that the pre-training dataset contains a wide variety of protein sequences, and—to a limited degree—is non-redundant. Having cluster or family metadata also makes it easier to sample stratified test and validation sets^8,10,41^.

Protein representation models may be evaluated internally, by analysing their representations, or externally, by benchmarking predictions based on the representations. It is sometimes interesting to visualize the embeddings generated by a protein representation model using a dimensionality reduction method, such as PCA or t-SNE^46^, to show that the representation model is able to capture biologically meaningful properties. Embeddings correlate with physicochemical properties^9,11,36,37^, the source organism proteome^9,11,36,37^, the secondary structure^11,36,37^, and the subcellular location^11,37^. The quality of these correlations generally improves with fine-tuning. Additionally, Vig et al.^47^ inspect the intermediate outputs of the TAPE Transformer model, and observe that, in the first layers, its attention heads specialize on amino acid type. Then, deeper layers focus on more complex features, such as binding sites and intra-chain protein contacts.

We observe a wide variety in the kinds of benchmarks used to evaluate these protein representation models. Even if the type of benchmark is similar, methods will frequently use different datasets, use different training and validation methodologies, and differ in how the datasets are preprocessed.

For the secondary structure prediction task, the model is asked to predict a structure (*α*-helix, *β*-strand, or coil) to regions of amino acids. There are two commonly used training sets, the NetsurfP-2.0 dataset^1^ and the SPOT dataset^48^. These datasets consists of training sequences with assigned secondary structure, derived from PDB structures. Another frequently used task is contact map prediction, which tests structural property prediction of a model^8,9,39,41,47,49^. For this task, a model needs to predict which residues are in close contact with each other within the tree-dimensional structure of a protein. Generally, contact maps need to be derived from protein structures and thus dataset are often based on the PDB.

Another type of benchmark task relates to whole protein labelling. The most common is remote homology detection. This is a classification task in which a model needs to predict if two sequences are homologous, i.e. descended during evolution from a common ancestor. Close homologs will be very similar in sequence, but protein sequence may diverge widely during evolution while function (and structure) remain conserved. The low sequence similarity of remote homologs makes this a difficult task. All articles we have evaluated use the SCOP database^50^ as the gold standard, which classifies protein domains into a hierarchical ontology. Pfam is also theoretically suitable for this, but the SCOP system attempts to classify into broader superfamilies than Pfam does, and thus is more suitable for remote—and more difficult to predict—homology.

Riesselman et al.^51^, Alley et al.^36^, and Rives et al.^9^ include variant effect prediction as a benchmark. In this task, the quantitative impact of a mutations is predicted for a specific set of protein functions, such as ligase activity and substrate binding. Two datasets are often used for this benchmark: firstly, the dataset used by Gray et al.^52^ includes 21 026 variant effect measurements of eight proteins from nine experimental datasets; secondly, Riesselman et al.^51^ combined 42 mutational scans, with a total of 712 218 mutations on 34 proteins.

Protein localization classification is also a protein-level labelling task. Protein function depends on subcellular localization. Abnormal localization can lead to dysfunction, which in turn can contribute to disease. Min et al.^40^, Heinzinger et al.^37^, and Elnaggar et al.^11^ use the DeepLoc dataset^53^, consisting of 13 858 proteins. Proteins are classified as being present in ten cellular locations, based on UniProt annotations. For transmembrane prediction, Bepler and Berger^39^, as well as Min et al.^40^ use the TOPCONS dataset^54^, containing 6 856 proteins. In this task, the model needs to predict for every amino acid in the training set whether it is membrane-spanning.

Notably, the Tape repository^8^ provides a set of 5 benchmark task for both biological properties prediction and protein engineering. For property prediction, the tasks included are generally also popular in the field for testing representation models.

Notable factors influencing protein representation model performance are the size of the training set, the model size, and pre-training using multiple (orthogonal) objectives. We find the CB513 dataset^55^, for secondary structure prediction on 8 classes (SS8), is commonly used to evaluate model quality, and is also sufficiently difficult to avoid ceiling effects; we summarize some of the best performing models on this dataset in Table 1. The state-of-the-art model NetSurfP-2.0^1^ achieves 72.3% accuracy on this set. From the protein representation models, only the MSA Transformer from Rao et al.^49^ achieves a higher accuracy: 72.9%. Big pre-trained Transformers come close to state-of-the-art performance: EMS-1b achieves 71.6%, ProtTrans-T5^11^ achieves 71.4%, and ProtTrans-Bert^11^ achieves 70% on SS8 prediction. This is in accordance with a general trend that we observe: most protein representation models do not outperform the state-of-the-art, which are often methods which are laboriously hand-tuned for optimum performance. However, large transformer-based models often come close on a wide variety of downstream tasks that it is debatable whether a statistically significant performance difference actually exists. Because the architectures are not specialized for a specific task—most are actually fairly straightforward conversions of natural language processing architectures—there is already significant value in being able to get to near state-of-the-art performance.

**Table 1.** Overview of the best performing protein representation models on the secondary structure prediction task for 8 classes (SS8).

| Model name | CB513 SS8 accuracy |  | Model size |  | Pre-training db size |  |
| --- | --- | --- | --- | --- | --- | --- |
| ProtBert-BFD | 70 | <sup>11</sup> | 420 M | <sup>11</sup> | 2 122 M | <sup>11</sup> |
| ProtTrans-T5 | 71.4±0.2 | <sup>49</sup> | 3 000 M | <sup>11</sup> | — | — |
| EMS-1b | 71.6±0.1 | <sup>49</sup> | 650 M | <sup>41</sup> | 250 M | <sup>41</sup> |
| EMS-MSA-1 | 72.9±0.2 | <sup>49</sup> | 100 M | <sup>49</sup> | 26 M | <sup>49</sup> |
| **NetSurfP-2.0 (mmseqs)** | **72.3** | <sup>1</sup> | — | — | no pretraining |  |
| **RaptorX** | **70.6** | <sup>1</sup> | — | — | no pretraining |  |

The accuracy in percentages is shown for the results on the CB513 dataset. Not all papers report significant numbers and standard deviations. Data from Klausen et al.<sup>1</sup> is shown in bold, as an example of state-of-the-art performance. Only the MSA Transformer (EMS-MSA-1) achieves higher accuracy compared to NetSurfP-2.0. The number of parameters (model size), the number of sequences in the pre-training dataset, and corresponding references are also reported.

Recently, Alphafold2^3^—a neural-network-based algorithm from Google’s DeepMind—gained a lot of attention for its protein structure prediction results at CASP-14. CASP is a bi-annual protein structure prediction competition^56^, where competitors are evaluated for their structure predictions on GDT, which is a score for structure similarity ranging from 0 to 100. Alphafold2 scored a medium GDT of 92.4, outcompeting other methods on most proteins and even leading to claims of solving the protein folding problem^3^. Alphafold2 takes as input a sequence, a MSA using the BFD database, and the 3D atom coordinates from homologs. In contrast with other methods we discuss here, Alphafold2 is not pre-trained.

## ProteinGLUE Benchmark tasks

The ProteinGLUE benchmark suite described in this work consists of the following seven benchmark tasks, which are all structural features that are labelled per amino acid in the protein sequence.

### Secondary structure (SS3 and SS8)

The dataset used for the secondary structure classification into three classes (*α*-helix, *β*-sheet, and coil) and into eight classes (coil, high-curvature, *β*-turn, *α*-helix, 310-helix, *π*-helix, *β*-strand, and *β*-bridge) was created by Hanson et al.^48^. This dataset is used in multiple prediction methods^2,48,57^. Proteins in the set were obtained using the PISCES server^58^, and were filtered on resolution <2.5Å, R-free <1 and sequence identity (seq.ID) 25%, according to BlastClust^42^. We split this dataset into 8 803 sequences for training, 1 102 sequences for validation and 1 102 sequences for testing. For the secondary structure prediction in three (SS3) and eight classes (SS8) these sets include the same proteins. Accuracy (ACC) is used for measuring model performance on these classification tasks.

### Solvent accessibility (ASA and BUR)

The dataset used for the solvent accessibility prediction is based on the same dataset used for SS3 and SS8. Training, validation and test sets were sampled independently from the secondary structure prediction sets, but include the same number of sequences for each. The absolute solvent accessibility (ASA) values, as given in the source data, were used to identify buried residues. Residues were determined as being buried (BUR)^59^ if the relative solvent accessible area—that is, the solvent accessible area divided by the maximal solvent accessible area for an amino acid type—was less than 7%^59^. Accuracy (ACC) is used for measuring model performance on the BUR classification task, and Pearson correlation coefficient (PCC) for the ASA regression task.

### Protein-protein interaction (PPI)

The dataset used for the PPI interface prediction task was created by Hou et al.^15^ for their random forest based PPI interface prediction model SeRenDIP^15,17^, which included both homodimer and heterodimer interfaces. The homomeric dataset was based on the Test_set 1 dataset of earlier work from Hou et al.^60^ and was filtered on 30% seq.ID. Remaining proteins were filtered at 25% seq.ID against the heterodimer dataset and the training sets of NetsurfP. The heteromeric dataset is based on the datasets Dset_186 and Dset_72 created by Murakami and Mizuguchi^61^, and were filtered at 25% seq.ID against the NetSurfP and DynaMine training, and the homomeric datasets^15^. We used those four datasets, retaining 287 homomeric training, 93 homomeric test, 118 heteromeric training and 44 heteromeric test proteins (the last stored protein for each was omitted). We then selected 20% of the homomeric and heteromeric training proteins individually in order to create a validation set. The homomeric and heteromeric training, validation and test sets respectively were concatenated into one training, validation and test set for the PPI task, containing both types of interfaces. The area under the receiver operating characteristic curve (AUC ROC) is used to evaluate model performance on the PPI prediction task.

### Epitope region (EPI)

The epitope dataset was obtained from Hou et al.^15^. This dataset is based on the structural antibody database for the Oxford Protein Information Group (SAbDab)^62^. The PDB structures of the antibody-antigen complexes were selected and antigen sequences were filtered at 25% seq.ID. We used the training, validation, and test sets of the first fold of the original 5-fold data split, which contains 179 training, 45 validation, and 56 test sequences. Area under the receiver operating characteristic curve (AUC ROC) is used for measuring model performance on the EPI prediction task.

### Hydrophobic patch prediction (HPR)

The hydrophobic patch dataset contains the structure-based assignment of hydrophobic patches, as created by Van Gils et al.^21^. PISCES was used to collect PDB structure of a resolution ≤ 3.0 Å, R-factor ≤ 0.3 and of sequence length between 40–10 000 residues. Only X-ray determined and non-C*α*-only structures were selected. Subsequently, the selected proteins were filtered on 25% seq.ID. Finally, only monomers were selected: transmembrane proteins were excluded. The resulting set contains 4 917 proteins. Because the model is unable to predict the total hydrophobic surface area, MolPatch^21^ was used to generate, per protein, the rank of each hydrophobic patch. For amino acids belonging to multiple patches the rank of the largest patch was assigned. The dataset was split into 60% training sequences, 15% validation sequences, and 25% test sequences. Pearson correlation coefficient (PCC) is used for measuring model performance on the HPR regression task.

Based on previous studies on protein structural properties using all kinds of prediction models, we expect SS3 and BUR to be *easy* prediction tasks followed by SS8 and ASA. The PPI and EPI prediction tasks are expected to be ‘*medium*’ difficult. The hydrophobic patch prediction is expected to be *hard* prediction task.

All the datasets of the ProteinGLUE benchmark suite are provided in the TensorFlow format, making the set easily reusable by the community. We refer to the section *TensorFlow format of datasets* in the supplement for detailed information about the data format.

## Pre-trained models

To set a challenging baseline for our datasets, we trained two transformer models based on the BERT architecture.^6^ We provide a `BERT medium` model, with 8 hidden layers, 8 attention heads and hidden size of 512, and we provide a `BERT base` model with 12 hidden layer, 12 attention heads and hidden size 768. The “hidden size” refers to the dimensionality of the vectors representing the tokens in the hidden layers (between transformer blocks).

### Pre-training data

Our baseline models were pre-trained on the Pfam dataset, more specifically the sequences from the PfamA 33.1 dataset^22^, filtered on 90% sequence identity. Similar to the BERT training process, we distinguish the amino acid sequences into regular sequences, which were at most 128 tokens long, and big ones at most 512.^6^ We discarded sequences longer than 512 amino acids, but as single domains longer than 512 amino acids are exceedingly rare, this does not exclude a significant fraction of the data. For both big and regular sequences, a test and a validation set were split off from the training data, each containing 10% of the total number of protein sequences. The resulting pre-training *training* dataset consists of 13 065 370 regular and 14 687 695 big sequences, a *validation* set of 1 469 855 regular and 1 835 963 big sequence, and a *test* set of 1 469 855 regular and 1 835 962 big sequences.

The downstream tasks may be used with models which have used other pre-training datasets than PfamA. In fact, in the natural language domain, progress in pre-training has often been the consequence of better-curated data, in addition to model improvements (in terms of size and architecture).^7^ We do, however, urge caution in assuming the source of performance improvements. If a new model and a different pre-training dataset are used, then, where possible, an ablation study should be performed.^63^ For this purpose we provide the precise, canonical subset of Pfam used for training our baseline models.

Unless mentioned otherwise, all aspects of the model were taken from the BERT model.^6^

### Pre-processing

Following the previously mentioned seperating of sequences into regular (max 128 tokens) and big (max 512 tokens), we tokenize sequences into amino acids, giving us a base vocabulary of 20. We also reserve 20 special tokens, used to annotate the sequence. Four of these, named PAD, CLS, MSK and SEP, are used in pre-training as explained below. The remaining 16 are reserved for potential use in downstream tasks. These are not used for any of our downstream tasks, but they may be useful for others.

While Devlin et al.^6^ slice fixed-length contiguous sub-sequences out of the corpus, we always train on full-length proteins. This means that our input sequences are variable length, so we pad each batch using PAD tokens so that all sequences within each batch have the same length.

### Pre-training methods

Following the structure of the original BERT models, with some slight deviations, we define two pre-training tasks:

#### Masked token prediction (MTP)

We change out a small percentage of tokens in the input sequences. The task is to reproduce the original tokens in the sequence. Some proportion of input tokens are *masked* (replaced by the masking token), and the others are *corrupted* by replacing them by randomly chosen, but different, amino acids; this is a change from the BERT setup because, with a vocabulary size of only 20 amino acids, the chance of randomising into the same amino acid gives a non-negligible performance boost. The model receives no indication which tokens are corrupted, but the loss is only computed over changed tokens. We use a 15% chance for a token to be changed. 80% are of these are masked, 10% are corrupted, and 10% to remain unchanged (but do contribute to the loss). Here we follow Devlin et al.^6^ precisely.

#### Next sentence prediction (NSP)

To stimulate the model to learn a representation of the whole sequence, a sequence-level task is added. That is, one label should be predicted for the whole sequence in addition to the ones for individual tokens. Devlin et al.^6^ created this task by either concatenating two half-length sequences from the corpus or using one whole-length sequence, and having the model predict which is the case. A special CLS token was prepended to every training example, and the representation of this token was used to predict the target label. We adapt the NSP task for the application to protein sequence data, by selecting either whole sequences from the data or concatenating the first part of one and the second part of another sequence chosen at random. Sequences are cut or left unchanged with equal probability. To make the NSP task more challenging, we cut each sequence randomly somewhere between 40-60% of its length to avoid having the cut be in the exact same position every time. We place a SEP token in between the two separate chunks of protein sequence, or in the middle of the complete sequence, using the same 40-60% selection to simulate a cutting point. Another SEP token is placed at the end, before the padding tokens. Following Devlin et al.^6^, we also add a learned segment embedding to every token which indicates whether the token belongs to the first or the second part of the sequence. Summed with the token- and position embeddings, these form the input embeddings.

#### Batching

For efficiency reasons, each batch either contains only regular sequences or only large sequences. We schedule the proportion of regular versus large sequence batches in each epoch. Training starts with 0% large sequence batches and linearly increases to 50% by the end of the run. Our assumption is that at the start of training, the model is not yet learning long dependencies, so it is more efficient to train on short sequences. Near the end of training, the model is hopefully learning longer dependencies, and the longer sequences become a valuable input.^†^

For each batch, we perform both tasks: the batch is made up of either original sequences or concatenated cut sequences, and in both cases, masking is applied as described above. We then compute losses for both the MTP and NSP tasks, using categorical cross-entropy, and add them together to produce the total loss.

### Pre-training results

Both the BERT `medium` and `base` model were pre-trained using the Layer-wise Adaptive Moments (LAMB) optimizer^64^. We used a learning rate of 0.00025 and batch sizes of 512 and 128 sequences for regular- and big batches, respectively. Both runs took approximately two weeks of wall-clock time of continuous training on four parallel TitanRTX GPUs and using mixed precision. The `medium` model was trained for 2 000 000 steps while the `base` one was trained for 1 000 000 steps.

Figure 1A shows the accuracy curves for masked token prediction (blue) and NSP (yellow) on training and validation data of the final pre-training run executed with a `medium` sized BERT model. This run was scheduled for 2 000 000 steps and converged on roughly 38% accuracy for the regular sequences in the masked token prediction task, and was still improving on the large sequences at the end of this run, as can be seen by the increasingly smaller downwards peaks in the bottom part of the plot. With longer training times, we would expect the accuracy for big sequences to also converge at around 38%, but the latest average training accuracy (light blue) after the full 2 000 000 steps at the end of this run was approximately 35%. The average validation (blue) and testing (dashed blue) accuracy for the masked token prediction were also both around 35% at the end. The NSP accuracy converged much faster than the masked token prediction and ended at an average training (light yellow), validation (yellow), and testing (dashed yellow) accuracy of around 96%.

**Figure 1.**
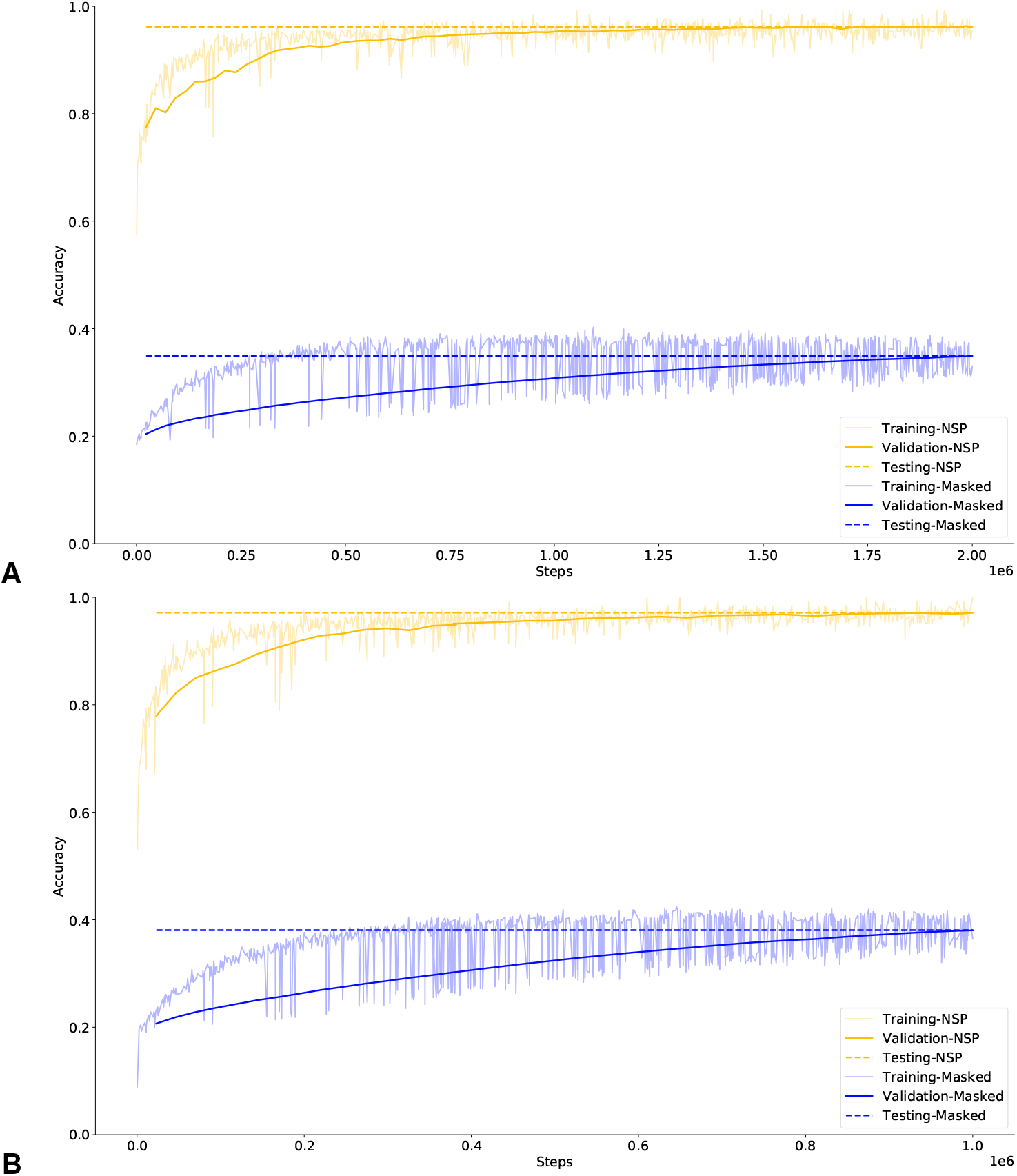
*Accuracy versus training steps* for masked token (blue) and next sentence (yellow) prediction on training (per batch) and validation (per epoch) data of (A) BERT `medium` and (B) BERT `base` pre-training runs. Dashed lines indicate the final testing data accuracy.

Figure 1B shows the same curves as panel A but for the BERT `base` model. As can be seen, the BERT `base` model shows a similar plot to the BERT `medium` model after only half the amount of steps. This increase in performance is likely due to its added complexity. The `base` model also performed better than the `medium` one overall, converging on roughly 41% accuracy for the regular sequences in the masked token prediction task, with the performance on big sequences still improving again, resulting in an average training (light blue) accuracy at the end of the 1 000 000 steps of 38%. The average validation (blue) and testing (dashed blue) accuracies at the end of this run were both approximately 38% as well. The NSP accuracy converged considerably faster again and ended at an average training (light yellow), validation (yellow), and testing (dashed yellow) accuracy of around 97%.

## Fine-tuned models

To provide baseline performance estimates, and as a proof-of-concept for pre-training performed as described in the previous section. We fine-tune both our BERT-derived models for all downstream tasks included in the ProteinGLUE benchmark dataset.

### Fine-tuning methods

#### Architecture and parameters

The fine-tuning models for the downstream tasks consists of the transformer model and a classifier. The size of the model is therefore dependent on the size of the pre-trained transformer. Classifiers of classification tasks (SS3, SS8, BU, PPI, and EPI) consist of a non-linear layer, a dropout layer, a classification layer, and a probability layer. The activation function in the non-linear layer is set to the Gaussian Error Linear Unit (GELU)^65^, which was also used in the pre-train layers. The dimension of the output space of the classification layer is set to the number of classes plus one, due to the padding of sequences. The activation function that is applied to the output in order to generate the probabilities is set to the softmax activation function. The classifier of the regression tasks (ASA and HPR) consisted of a linear layer, a dropout layer, and a regression layer. The activation function of the linear layer is also set to the GELU activation function. The dimensionality of the output space in the regression layer is set to 1.

We tuned the hyperparameters *batch size, learning rate* and *dropout rate* for all downstream tasks by training models on the training sets and compare performances on the validation set. We performed an exhaustive grid search on the pre-trained `medium` and `base` models separately. The dropout rate is the rate included in the dropout layer of the fine-tuning classifier. For the `medium` model we considered batch sizes 8, 16 and 32, learning rates 6.25e-5, 1.25e-4, 2.5e-4 and dropout rates 0.0, 0.05, 0.1 and 0.2. When a high performance is attained for the largest learning rate, the learning rate is further increased to 5.0e-4 or 1.0e-3. For the `base` model we considered the same values, except for the batch size where only batch sizes of 4 and 8 were included due to memory limitations. After the exhaustive grid search, the 4 to 8 best performing hyper-parameter sets were selected for each prediction task. The models were trained 4 times on each set after which the best hyper-parameters were selected, based on the mean performance (see supplementary Table 1).

We decided to set the maximum length of the considered sequences to 512 and keep this value constant over all downstream prediction tasks. Even as for the pre-training, the LAMB optimiser was used. We define the fine-tuning models to be converged when they are trained for 15 epochs or 2 000 steps.

For the protein interface and epitope prediction tasks, we had to deal with the class imbalance of the dataset. The PPI interface training set consisted of 16 605 residues indicated as interface residues, and 66 180 residues indicated as non-interacting. The epitope training set consisted of 4 503 and 41 938 residues indicated as interacting and non-interacting, respectively. We included the ratio of the number of non-interface residues over the number of interface residues as weight in the loss function. Therefore this weight was set to 3.99 for the PPI interface prediction and 9.31 for the epitope prediction.

All downstream tasks were trained and validated on a single compute node, consisting of a single TitanX GPU containing 12Gb of GPU memory, with a (wallclock) run time of about 4 hours.

#### Training and evaluation

During training, the loss was determined by categorical cross entropy for classification tasks, and mean absolute error for regression tasks. Model performance was assessed by the Pearson correlation coefficient (PCC) for the regression tasks (ASA and HPR), accuracy (ACC) for the classification of the structural components (SS3 and SS8) and the identification of the buried residues, and the area under the receiver operating characteristic curve (AUC ROC) for the interface predictions (PPI and EPI). Note that the range of the PCC is between −1 and 1, whereas the range of the ACC and AUC is between 0 and 1. We determined random prediction performances for all downstream prediction tasks in the benchmark set. For the SS3, SS8, and BUR prediction task this random performance is set to the fraction of the number of majority-class residues over all residues. For the ASA and HPR we compare the labels with a sequence of the same size sampled from a standard normal distribution. The expected random expected performance, in this way, is close to zero. For the two interface prediction tasks we set the baseline to the random performance of a ROC curve, which is 0.5.

The previously described pre-trained `medium` and `base` models were used for the hyperparameter tuning. The performance, per downstream task, between the `medium` and `base` model were compared. For comparison, we also trained the downstream tasks on both excluding the pre-training step. During training of the pre-training models two checkpoints were stored manually. This includes the checkpoints of the `medium` model at step 500 000 and 1 600 000, and of the `base` model at step 350 00 and 700 000. The predictive performance of the different downstream tasks was evaluated over these checkpoints to check for overfitting. All fine-tuning models were trained ten times, after which the mean performance and standard error on the validation set was determined.

### Fine-tuning results

In this study, we selected seven protein structural prediction tasks to evaluate pre-trained transformer models in a varied way. Our pre-trained models include a BERT `medium` and a BERT `base` model trained on protein domain sequences from the PfamA database. The datasets for these tasks were based on previous state-of-the-art prediction studies on these tasks. Each selected dataset was divided into a training, validation and test set, see Table 2.

**Table 2.**
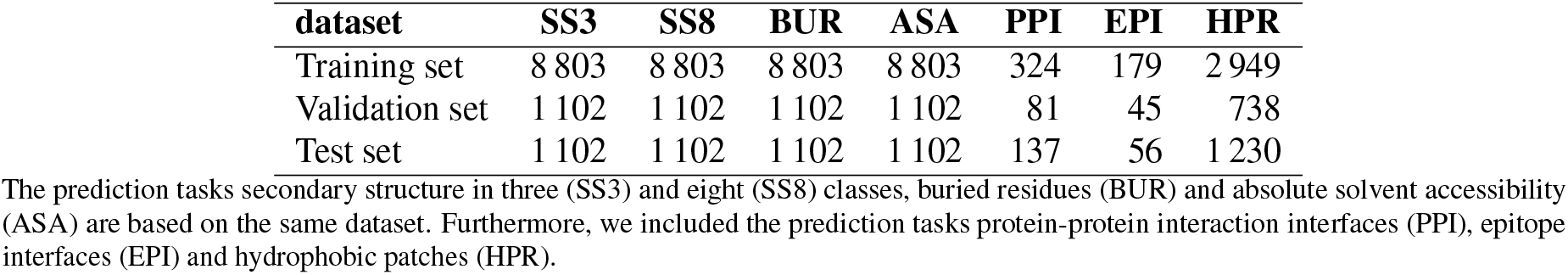
Overview of the number of protein sequences in the training, validation and test set of the seven downstream protein structural prediction tasks.

#### Hyperparameter tuning

Similar to the study of Devlin et al.^6^, most hyperparameters were set to the parameters of the pre-training-tasks. However, the downstream tasks are considered as having converged after reaching 2 000 steps, or 15 epochs. We tuned the the batch size, learning rate and dropout rate for each task specific on both the converged `medium` model and the converged `base` model, see Table 3.

**Table 3.** The tuned hyperparameters for the `medium` and `base` models.

| Model | Pre-training task | Batch size | Learning rate | Dropout rate |
| --- | --- | --- | --- | --- |
| medium | SS3 | 16 | 5e-4 | 0.1 |
|  | SS8 | 16 | 5e-4 | 0.05 |
|  | BUR | 8 | 5e-4 | 0.05 |
|  | ASA | 16 | 2.5e-4 | 0.05 |
|  | PPI | 8 | 1.25e-4 | 0.05 |
|  | EPI | 16 | 2.5e-4 | 0.05 |
|  | HPR | 32 | 6.25e-5 | 0.05 |
| base | SS3 | 8 | 2.5e-4 | 0.0 |
|  | SS8 | 8 | 2.5e-4 | 0.0 |
|  | BUR | 8 | 2.5e-4 | 0.1 |
|  | ASA | 8 | 1.25e-4 | 0.0 |
|  | PPI | 8 | 6.25e-5 | 0.0 |
|  | EPI | 8 | 1.25e-4 | 0.1 |
|  | HPR | 8 | 6.25e-5 | 0.1 |

The hyperparameters are tuned for each downstream structural prediction task individually on both the converged `medium` pre-trained model and the converged `base` pre-trained model using the validation sets.

#### Model performance

The pre-trained models were used as a basis to train and evaluate models for the seven fully supervised downstream tasks. It is commonly assumed that, by allowing the model to learn the structure of the input data on a large amount of unlabeled sequences, the performance on specific tasks can be improved, without requiring large amounts of labeled sequences, which are often difficult or expensive to acquire. Figure 2 tests this assumption using our data and models. We compare the performance of a `base` and `medium` model on the downstream tasks, with and without pre-training. The results show a clear improvement of pre-training for 6 out of the 7 downstream tasks. For the epitope prediction both the `medium` and `base` pre-trained models perform slightly worse compared to the non pre-trained models. On the validation set, the improvement of pre-training is shown for all tasks (see Supplementary Figure 1). For the hydrophobic patch regression (HPR), however, the results are less clear. There is minimal improvement in the `medium` model. The `base` model does show improvement, but that is because the model without pre-training far under performs, compared to the `medium` version. This appears to be the most challenging task in our set of benchmarks, and the one for which models behave the most counter-intuitively.

**Figure 2.**
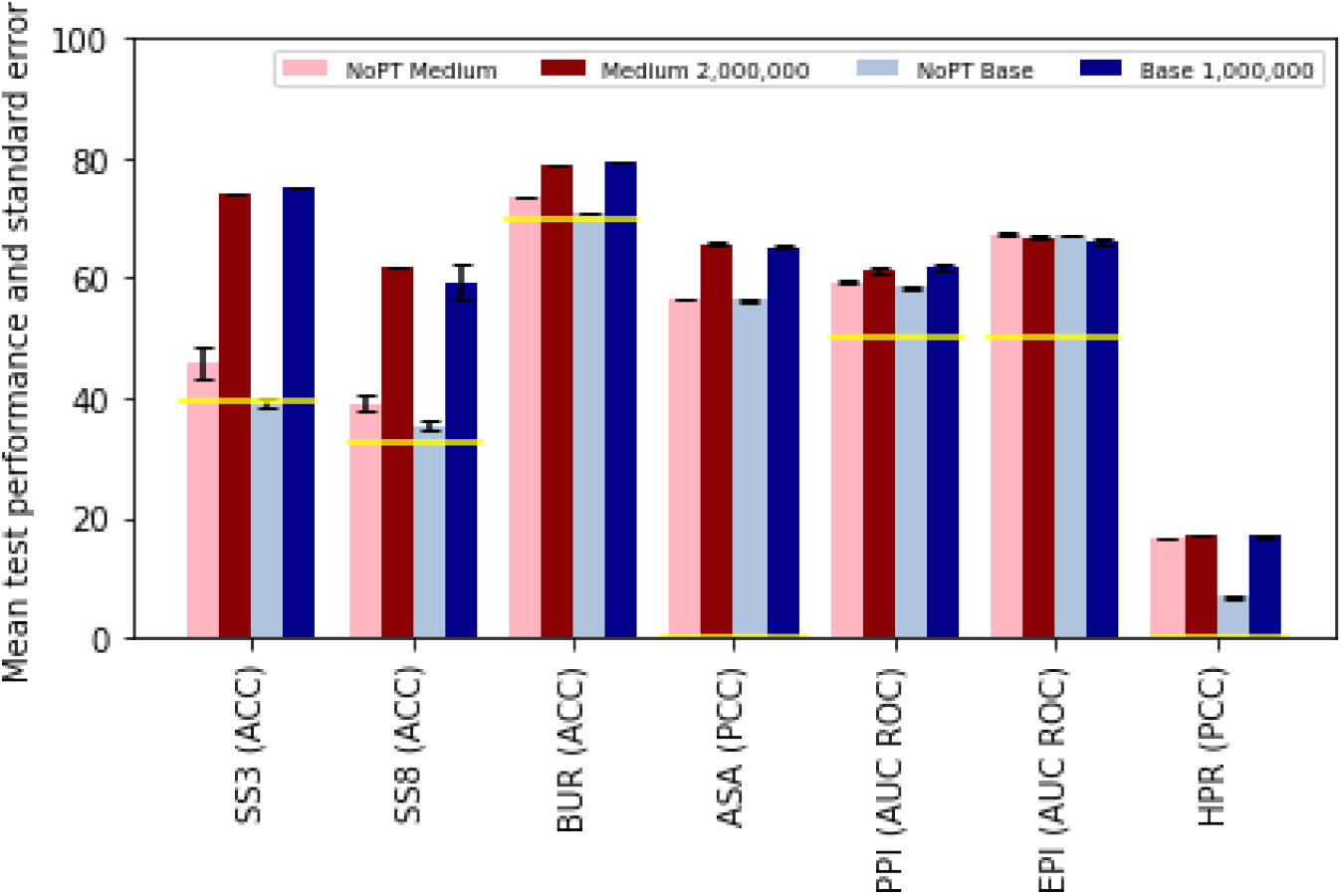
Improved performance on downstream tasks with pre-trained models. Prediction performances of the benchmark test set, as introduced in section *ProteinGLUE Benchmark tasks*, for a `medium` model without the pre-training step (pink), `medium` model including a pre-trained model until step 2 000 000 (dark red), `base` model without the pre-training step (grey), and `base` model including a pre-trained model until step 1 000 000 (dark blue). The yellow lines indicate the performance of a random or majority-class baseline. All models are trained ten times on their selected set of hyperparameters after which the mean performance and standard error is determined.

Furthermore, we compare the downstream task performances against their random performance. The random performance of the SS3, SS8 and BUR prediction was set to the majority-class which resulted in 39%, 32%, and 70% respectively. The random performance of the ASA prediction was estimated to be -8.9e-4, and of the HPR prediction to -6.0e-4, i.e. both very close to zero. The random performance of the AUC ROC performance measure was set to the value 0.5. We conclude that in all cases except for the not pre-trained `base` model on the SS8 and BUR prediction tasks, the model performances, including standard error bars, outperform the random performances.

To check for convergence in the pre-trained models we also monitored the performance of the downstream task during pre-training, which is shown in Figure 3. Results on the validation set are shown in Supplementary Figure 2. Our main observation here is that for most tasks, pre-training confers a strong advantage. For the `medium` model (Figure 3A), the performance increases with the amount of pre-training, aside from some small fluctuations. For the larger `base` model (Figure 3B), we note a more uneven progress in the number of training steps, with greater standard error.

**Figure 3.**
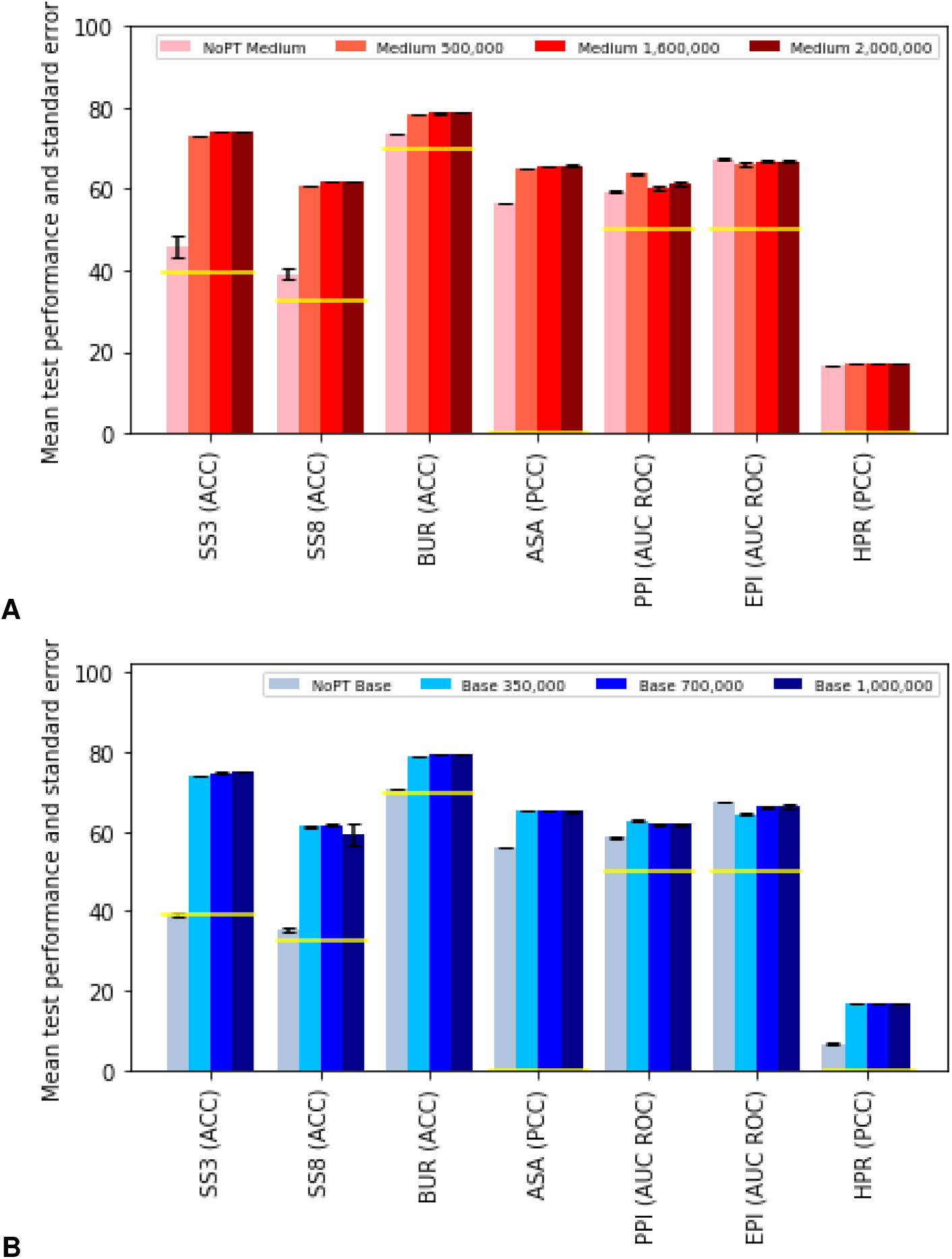
Performance of downstream tasks monitored during pre-training. Prediction performances of the benchmark test set, as introduced in section *ProteinGLUE Benchmark tasks*, (A) for a `medium` sized model, and (B) for a `base` sized model, trained without the pre-training step (pink/grey), and on a pre-trained model for 350 000 (light red/blue), 700 000 (red/blue) and 1 000 000 steps (dark red/blue). Further details as in Figure 2.

Figure 4 shows the difference between the validation and test performance. For some tasks the validation performance is substantially higher than the test performance. To some degree this is to be expected, but it may also indicate over-tuning of hyperparameters. Overall, however, model performance is stable over the test and validation sets.

**Figure 4.**
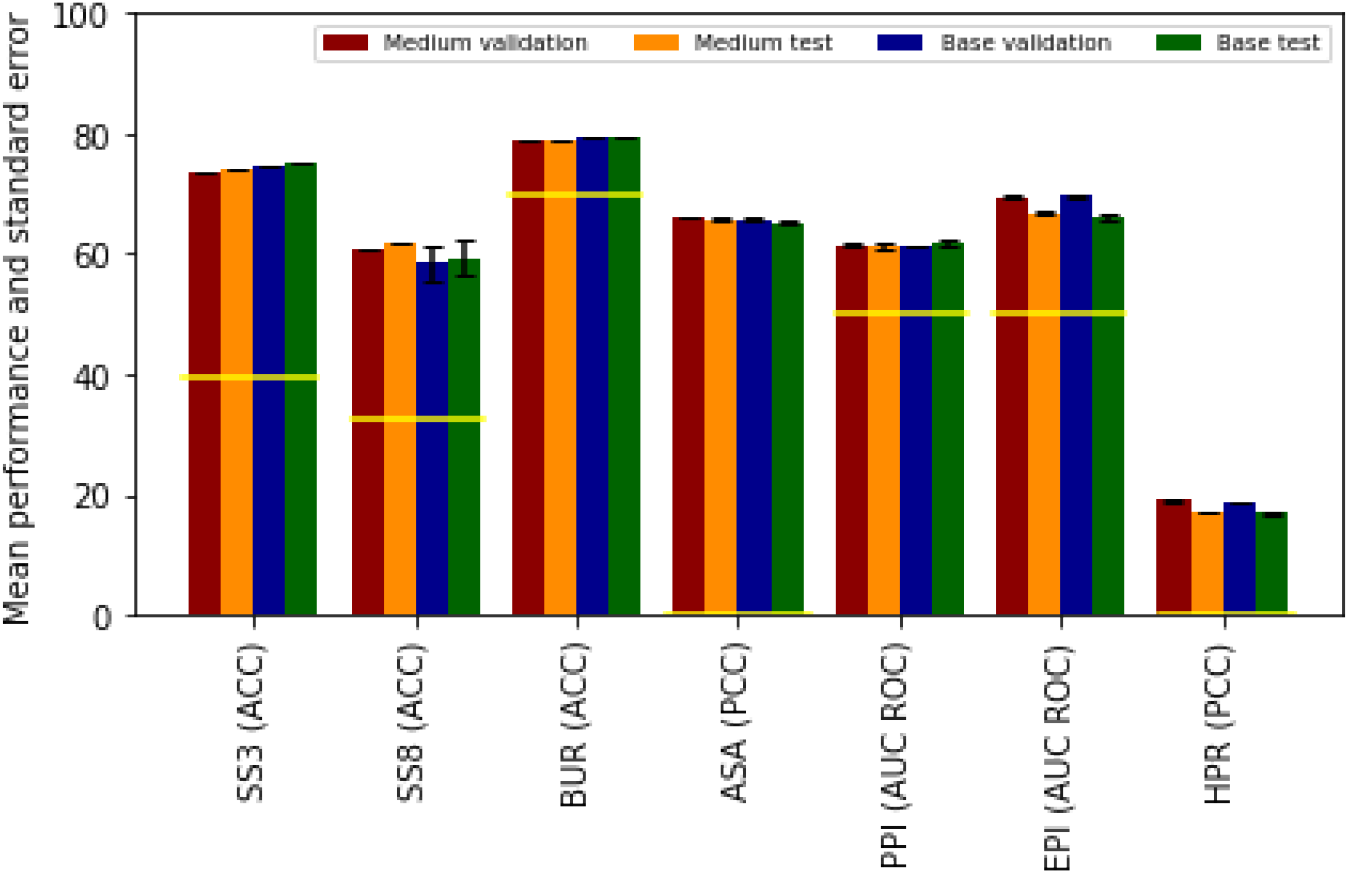
Performance on downstream task is generally stable between test and validation sets. Prediction performances of the benchmark validation and test set for both the converged `medium` and `base` models. Further details are given in Figure 2.

## Discussion

We present the ProteinGLUE benchmark set: a collection of classification and regression tasks in the protein sequence domain. Our main aim with this resource is to provide a standardized way to evaluate pre-trained protein models, and to provide clearer and more informative comparison between such models. We have also provided initial results of baseline models and checkpoints of these models for further development. All code used to train our models is available online at https://github.com/ibivu/protein-glue.

We pre-trained two transformer models inspired by the BERT `medium` and `base` architectures. Pre-training was performed with two objectives commonly used in NLP: masked symbol prediction; and next sentence prediction, which we adapted to predict matching halves of a protein sequence. Given the fast convergence on the NSP objective—with a final accuracy close to 100% (Figure 1)—in future work it could be investigated how this task could be made more difficult and how a harder pre-training task may help improve downstream performance. We evaluated prediction performances on the ProteinGLUE benchmark set on secondary structure in three (SS3) and eight (SS8) classes, buried residues (BUR) and absolute solvent accessibility (ASA), protein-protein interaction interfaces (PPI), epitope interfaces (EPI) and hydrophobic patches (HPR) using the `medium` and `base` pre-trained BERT models. We compared results against performances of these models without pre-training (Figure 2). Except for the epitope interface (EPI) prediction, all other benchmark tasks (SS3, SS8, BUR, ASA, PPI, HPR) achieved better performance on the pre-trained models. As expected the hydrophobic patch (HPR) prediction is the most challenging task.

The larger `base` model does not always outperform the `medium` model (Figure 2). However, the differences are small and not statistically significant, and are not indicative of a lack of convergence for the `base` model. Additionally, for some datasets, performance does not increase monotonically with the number of training steps (Figure 3). While these instances are rare, it suggests there may be some benefit to early stopping in pre-training, or added regularization to allow for a more uniform convergence. We leave this as a matter for future work.

While it is not our aim to outperform state of the art structural prediction methods, which typically use feature sets that were handcrafted and tuned over many years of painstaking research (e.g. Table 1 for SS8), we compare the ProteinGLUE benchmark set results against these methods in order to provide a general understanding of the performances. Note that for most comparisons the test sets are different, so the observed differences in performance should only be taken as a rough approximation. The OPUS-TASS method^2^ outperforms earlier studies on the secondary structure prediction based on the dataset created by Hanson et al.^48^, which is also used here. OPUS-TASS reaches 89% and 79% on SS3 and SS8 prediction, respectively, on one of their test sets. We reach 75% accuracy on the pre-trained `base` model for SS3 and 62% accuracy on the pre-trained `medium` model for SS8. The SeRenDIP method^15,17^, trained on the combined dataset of homodimer and heterodimer protein-protein interactions, resulted in an AUC ROC of 0.72 on the homodimer test set and 0.64 on the heterodimer test set. We reach an AUC ROC for PPI of 0.62 on the combined test set. The SeRenDIP-CE method^19^ for epitope interface prediction reaches an average AUC ROC of 0.69 over their 5 folds. We reached an average AUC ROC for EPI of 0.67 over ten times training the first fold.

Compared to the standardized benchmarks available in the NLP domain, a set of seven tasks is a modest start. We hope that further benchmark sets will follow ours. All tasks included in our ProteinGLUE benchmark set currently label individual amino acids: no tasks were included for which the label applies to the whole sequence, or to a region such as a protein domain. Including tasks like fold or function prediction may provide such opportunities. We expect our ProteinGLUE benchmark set will prove to be useful in itself, and that combined with the baseline models and performance comparison presented here, it will provide a starting point for further improvement of deep learning approaches, and transformer-based models in particular, to the exciting field of protein structural and functional property prediction.

## Acknowledgements

H.C. and R.W. would like to thank The Network Institute VU for funding this project. This work was carried out on the Dutch national e-infrastructure with the support of SURF Cooperative and on the distributed ASCII supercomputer^66^.

## Author contributions statement

M.D. conceived the prototype model and implemented initial experiments. H.C., R.W. implemented final versions of all code, and performed final experiments, under supervision from M.D., P.B. and K.A.F. R.V. Performed an extensive literature review, summarized here. All authors contributed to writing and editing of the manuscript.

## Footnotes

* Code: https://github.com/ibivu/protein-glue

* https://www.tensorflow.org/api_docs/python/tf/io/TFRecordWriter

† The memory use of transformer models grows quadratically in the sequence length, so a batch of many short sequences is more efficient to process than a batch of few long sequences, even if the total number of tokens is equal.

## References

1. Klausen, M. S. et al. Netsurfp-2.0: Improved prediction of protein structural features by integrated deep learning. Proteins: Struct. Funct. Bioinforma. 87, 520–527 (2019).

2. Xu, G., Wang, Q. & Ma, J. OPUS-TASS: a protein backbone torsion angles and secondary structure predictor based on ensemble neural networks. Bioinformatics 36, 5021–5026, DOI: 10.1093/bioinformatics/btaa629 (2020). https://academic.oup.com/bioinformatics/article-pdf/36/20/5021/35065215/btaa629.pdf.

3. Jumper, J. et al. Highly accurate protein structure prediction with alphafold. Nature 596, 583–589 (2021).

4. Sejnowski, T. J. The unreasonable effectiveness of deep learning in artificial intelligence. Proc. Natl. Acad. Sci. 117, 30033–30038 (2020).

5. Liu, X., He, P., Chen, W. & Gao, J. Improving multi-task deep neural networks via knowledge distillation for natural language understanding. arXiv preprint arXiv:1904.09482 (2019).

6. Devlin, J., Chang, M.-W., Lee, K. & Toutanova, K. Bert: Pre-training of deep bidirectional transformers for language understanding. arXiv preprint arXiv:1810.04805 (2018).

7. Radford, A. et al. Language models are unsupervised multitask learners. OpenAI blog 1, 9 (2019).

8. Rao, R. et al. Evaluating protein transfer learning with tape. Adv. neural information processing systems 32, 9689 (2019).

9. Rives, A. et al. Biological structure and function emerge from scaling unsupervised learning to 250 million protein sequences. Proc. Natl. Acad. Sci. 118 (2021).

10. Madani, A. et al. Progen: Language modeling for protein generation. arXiv preprint 2004.03497 (2020).

11. Elnaggar, A. et al. Prottrans: towards cracking the language of life’s code through self-supervised deep learning and high performance computing. arXiv preprint 2007.06225 (2020).

12. Wang, A. et al. Glue: A multi-task benchmark and analysis platform for natural language understanding. arXiv preprint 1804.07461 (2018).

13. Pauling, L., Corey, R. B. & Branson, H. R. The structure of proteins: two hydrogen-bonded helical configurations of the polypeptide chain. Proc. Natl. Acad. Sci. 37, 205–211 (1951).

14. Kabsch, W. & Sander, C. Dictionary of Protein Secondary Structure: Pattern Recognition of Hydrogen-Bonded and Geometrical Features. Biopolymers 22, 2577–2637 (1983).

15. Hou, Q., Geest, P., Vranken, W. & Feenstra, K. A. Seeing the trees through the forest: Sequence-based homo-and heteromeric protein-protein interaction sites prediction using random forest. Bioinformatics 33, 1479–1487, DOI: 10.1093/bioinformatics/btx005 (2017).

16. Bork, P. et al. Predicting Function: From Genes to Genomes and Back. J. Mol. Biol. 283, 707–725, DOI: 10.1006/jmbi.1998.2144 (1998).

17. Hou, Q. et al. SeRenDIP: SEquential REmasteriNg to DerIve profiles for fast and accurate predictions of PPI interface positions. Bioinformatics 35, 4794–4796, DOI: 10.1093/bioinformatics/btz428 (2019). https://academic.oup.com/bioinformatics/article-pdf/35/22/4794/30706825/btz428.pdf.

18. Potocnakova, L., Bhide, M. & Pulzova, L. B. An introduction to b-cell epitope mapping and in silico epitope prediction. J. immunology research 2016 (2016).

19. Hou, Q. et al. Serendip-ce: sequence-based interface prediction for conformational epitopes. Bioinformatics 37, 3421–3427 (2021).

20. Dill, K. A. Theory for the folding and stability of globular proteins. Biochemistry 24, 1501–1509 (1985).

21. Gils, J. et al. How sticky are our proteins?: Quantifying hydrophobicity of the human proteome. arXiv.org (2021).

22. Mistry, J. et al. Pfam: The protein families database in 2021. Nucleic Acids Res. 49, D412–D419 (2021).

23. Sajjad, H., Dalvi, F., Durrani, N. & Nakov, P. On the effect of dropping layers of pre-trained transformer models. arXiv preprint 2004.03844 (2020).

24. Nugroho, K. S., Sukmadewa, A. Y. & Yudistira, N. Large-scale news classification using bert language model: Spark nlp approach. In 6th International Conference on Sustainable Information Engineering and Technology 2021, 240–246 (2021).

25. Blum, A. & Mitchell, T. Combining labeled and unlabeled data with co-training. In Proceedings of the eleventh annual conference on Computational learning theory, 92–100 (1998).

26. Chapelle, O., Sch olkopf, B. & Zien, A. Semi-supervised learning (chapelle, o. et al., eds.; 2006)[book reviews]. IEEE Transactions on Neural Networks 20, 542–542 (2009).

27. Scudder, H. Probability of error of some adaptive pattern-recognition machines. IEEE Transactions on Inf. Theory 11, 363–371, DOI: 10.1109/TIT.1965.1053799 (1965).

28. Howard, J. & Ruder, S. Universal language model fine-tuning for text classification. arXiv preprint 1801.06146 (2018).

29. Peters, M. E. et al. Deep contextualized word representations. In Proceedings of NAACL-HLT, 2227–2237 (2018).

30. Stringer, B. et al. Pipenn: Protein interface prediction with an ensemble of neural nets. bioRxiv DOI: 10.1101/2021.09.03.458832 (2021).

31. Vaswani, A. et al. Attention is all you need. In Advances in neural information processing systems, 5998–6008 (2017).

32. Brown, T. B. et al. Language models are few-shot learners. In Larochelle, H., Ranzato, M., Hadsell, R., Balcan, M. & Lin, H. (eds.) Advances in Neural Information Processing Systems 33: Annual Conference on Neural Information Processing Systems 2020, NeurIPS 2020, December 6-12, 2020, virtual (2020).

33. Yang, Y. et al. Sixty-five years of the long march in protein secondary structure prediction: the final stretch? Briefings bioinformatics 19, 482–494 (2018).

34. Chen, L. et al. Transformercpi: improving compound–protein interaction prediction by sequence-based deep learning with self-attention mechanism and label reversal experiments. Bioinformatics 36, 4406–4414 (2020).

35. Yao, Y., Du, X., Diao, Y. & Zhu, H. An integration of deep learning with feature embedding for protein– protein interaction prediction. PeerJ 7, e7126 (2019).

36. Alley, E. C., Khimulya, G., Biswas, S., AlQuraishi, M. & Church, G. M. Unified rational protein engineering with sequence-based deep representation learning. Nat. methods 16, 1315–1322 (2019).

37. Heinzinger, M. et al. Modeling aspects of the language of life through transfer-learning protein sequences. BMC bioinformatics 20, 1–17 (2019).

38. Strodthoff, N., Wagner, P., Wenzel, M. & Samek, W. Udsmprot: universal deep sequence models for protein classification. Bioinformatics 36, 2401–2409 (2020).

39. Bepler, T. & Berger, B. Learning protein sequence embeddings using information from structure. arXiv preprint 1902.08661 (2019).

40. Min, S. et al. Pre-training of deep bidirectional protein sequence representations with structural information. IEEE Access 9, 123912–123926 (2021).

41. Rives, A. et al. Biological structure and function emerge from scaling unsupervised learning to 250 million protein sequences. Proc. Natl. Acad. Sci. 118 (2021).

42. Altschul, S. F. et al. Gapped blast and psi-blast: a new generation of protein database search programs. Nucleic acids research 25, 3389–3402 (1997).

43. Uniprot: the universal protein knowledgebase in 2021. Nucleic Acids Res. 49, D480–D489 (2021).

44. Steinegger, M. & Söding, J. Clustering huge protein sequence sets in linear time. Nat. communications 9, 1–8 (2018).

45. Steinegger, M., Mirdita, M. & Söding, J. Protein-level assembly increases protein sequence recovery from metagenomic samples manyfold. Nat. methods 16, 603–606 (2019).

46. Van der Maaten, L. & Hinton, G. Visualizing data using t-sne. J. machine learning research 9 (2008).

47. Vig, J. et al. Bertology meets biology: Interpreting attention in protein language models. arXiv preprint 2006.15222 (2020).

48. Hanson, J., Paliwal, K., Litfin, T., Yang, Y. & Zhou, Y. Accurate prediction of protein contact maps by coupling residual two-dimensional bidirectional long short-term memory with convolutional neural networks. Bioinformatics 34, 4039–4045 (2018).

49. Rao, R. et al. Msa transformer. bioRxiv DOI: 10.1101/2021.02.12.430858 (2021). https://www.biorxiv.org/content/early/2021/02/13/2021.02.12.430858.full.pdf.

50. Fox, N. K., Brenner, S. E. & Chandonia, J.-M. Scope: Structural classification of proteins—extended, integrating scop and astral data and classification of new structures. Nucleic acids research 42, D304–D309 (2014).

51. Riesselman, A. J., Ingraham, J. B. & Marks, D. S. Deep generative models of genetic variation capture the effects of mutations. Nat. methods 15, 816–822 (2018).

52. Gray, V. E., Hause, R. J., Luebeck, J., Shendure, J. & Fowler, D. M. Quantitative missense variant effect prediction using large-scale mutagenesis data. Cell systems 6, 116–124 (2018).

53. Almagro Armenteros, J. J., Sønderby, C. K., Sønderby, S. K., Nielsen, H. & Winther, O. Deeploc: prediction of protein subcellular localization using deep learning. Bioinformatics 33, 3387–3395 (2017).

54. Tsirigos, K. D., Peters, C., Shu, N., Käll, L. & Elofsson, A. The topcons web server for consensus prediction of membrane protein topology and signal peptides. Nucleic acids research 43, W401–W407 (2015).

55. Cuff, J. A. & Barton, G. J. Evaluation and improvement of multiple sequence methods for protein secondary structure prediction. Proteins: Struct. Funct. Bioinforma. 34, 508–519 (1999).

56. Kryshtafovych, A., Schwede, T., Topf, M., Fidelis, K. & Moult, J. Critical assessment of methods of protein structure prediction (casp)—round xiii. Proteins: Struct. Funct. Bioinforma. 87, 1011–1020 (2019).

57. Hanson, J., Paliwal, K., Litfin, T., Yang, Y. & Zhou, Y. Improving prediction of protein secondary structure, backbone angles, solvent accessibility and contact numbers by using predicted contact maps and an ensemble of recurrent and residual convolutional neural networks. Bioinforma. (Oxford, England) 35, 2403–2410, DOI: 10.1093/bioinformatics/bty1006 (2019).

58. Wang, G. & Dunbrack Jr, R. L. Pisces: a protein sequence culling server. Bioinformatics 19, 1589–1591 (2003).

59. Hubbard, T. & Blundell, T. Comparison of solvent-inaccessible cores of homologous proteins: definitions useful for protein modelling. Protein Eng. Des. Sel. 1, 159–171 (1987).

60. Hou, Q., Dutilh, B. E., Huynen, M. A., Heringa, J. & Feenstra, K. A. Sequence specificity between interacting and non-interacting homologs identifies interface residues–a homodimer and monomer use case. BMC bioinformatics 16, 1–12 (2015).

61. Murakami, Y. & Mizuguchi, K. Applying the naïve bayes classifier with kernel density estimation to the prediction of protein–protein interaction sites. Bioinformatics 26, 1841–1848 (2010).

62. Dunbar, J. et al. Sabdab: the structural antibody database. Nucleic acids research 42, D1140–D1146 (2014).

63. Lipton, Z. C. & Steinhardt, J. Research for practice: troubling trends in machine-learning scholarship. Commun. ACM 62, 45–53 (2019).

64. You, Y. et al. Large batch optimization for deep learning: Training bert in 76 minutes. arXiv preprint 1904.00962 (2019).

65. Hendrycks, D. & Gimpel, K. Bridging nonlinearities and stochastic regularizers with gaussian error linear units. CoRR abs/1606.08415 (2016). 1606.08415.

66. Bal, H. et al. A medium-scale distributed system for computer science research: Infrastructure for the long term. Computer 49, 54–63 (2016).

