## Supplementary information, figures and tables for "ProteinGLUE: A multi-task benchmark suite for self-supervised protein modeling"

### Supplementary information for ProteinGLUE: A multi-task benchmark suitefor self-supervised protein modeling.

[1] Vrije Universiteit, Informatics Institute, Amsterdam, 1081 HV, Netherlands
[+] these authors contributed equally to this work

#### Technical implementation details

##### Tensorflow format of datasets

The protein sequences and their corresponding labels were extracted from all the fine-tuning data sets and converted to a `TFRecord` file using `TFRecordWriter`.[1] The `TFrecord` format stores sequences and binary records. During the transformation the `tf.train.Example` message is used for the `""string": value mapping"`. This mapping can except the feature types bytes, float and int64. The protein sequence is therefore converted to a bytes list. The absolute solvent accessibility values is converted to a float. All other prediction tasks are stored as integers. The fine-tuning tasks SS3, SS8, BUR, PPI and EPI are binary values. For the hydrophobic patch prediction the patches were ranked per protein based on the size of the patch. Therefore, the patch value was indicated as integer.

##### Negative results

We tried using a reduced target alphabet of only 12 tokens for some of our experiments within the pre-training procedure because there is a lot of variation in the sequences possible for a protein with a given shape[?,?]. Hence, we tested whether mapping these similar amino acids to the same tokens would improve model performance since we did not want to penalize the model for predicting an amino acid which is largely the same in a protein structural context as the strictly correct one in that specific sequence. However, what we found was that even though using a reduced target alphabet obviously increased the performance metrics on the pre-training task, we did not observe a notable difference in performance on the downstream tasks.

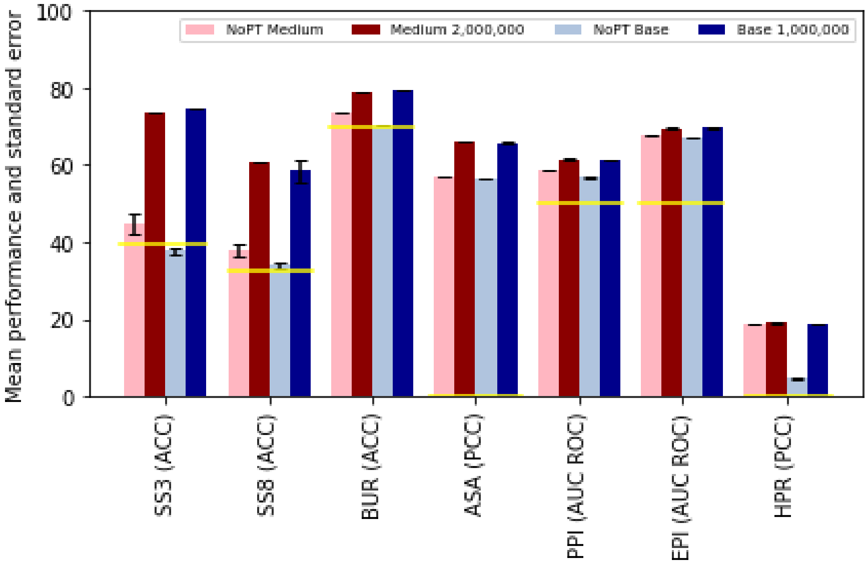


**Figure 1.** *Improved performance on fine-tuning tasks with pre-trained models.* Prediction performances of the benchmark validation set, for a `medium` model without the pre-training step (pink), `medium` model including a pre-trained model until step 2 000 000 (dark red), `base` model without the pre-training step (grey), and `base` model including a pre-trained model until step 1 000 000 (dark blue). The yellow lines indicate the performance of a random or majority-class baseline. All models are trained ten times on their selected set of hyperparameters after which the mean performance and standard error is determined.

**Table 1.** *The results of tuning the hyperparameters for the converged* `medium` *and* `base` *models.*

| Model | Pre-training task | Batch size | Learning rate | Dropout rate | mean performance |
|---|---|---|---|---|---|
| `medium` | SS3 | 16 | 2.5e-4 | 0.0 | 73.28 ACC |
| | | 16 | 2.5e-4 | 0.05 | 73.43 ACC |
| | | 16 | 5.0e-4 | 0.05 | 73.50 ACC |
| | | 32 | 2.5e-4 | 0.05 | 73.30 ACC |
| | | 16 | 5.0e-4 | 0.1 | **73.57** ACC |
| | | 32 | 2.5e-4 | 0.1 | 73.33 ACC |
| | | 32 | 5.0e-4 | 0.05 | 73.36 ACC |
| | | 8 | 5.0e-4 | 0.05 | 73.52 ACC |
| | SS8 | 8 | 5.0e-4 | 0.05 | 53.66 ACC |
| | | 16 | 5.0e-4 | 0.05 | **60.98** ACC |
| | | 32 | 2.5e-4 | 0.05 | 60.52 ACC |
| | | 32 | 5.0e-4 | 0.05 | 60.62 ACC |
| | | 16 | 5.0e-4 | 0.1 | 60.70 ACC |
| | | 32 | 5.0e-4 | 0.1 | 60.83 ACC |
| | BUR | 8 | 2.5e-4 | 0.0 | 78.75 ACC |
| | | 16 | 2.5e-4 | 0.0 | 78.77 ACC |
| | | 8 | 5.0e-4 | 0.05 | **78.92** ACC |
| | | 16 | 2.5e-4 | 0.05 | 78.88 ACC |
| | | 16 | 2.5e-4 | 0.1 | 78.92 ACC |
| | | 32 | 2.5e-4 | 0.05 | 78.78 ACC |
| | | 16 | 5.0e-4 | 0.1 | 78.81 ACC |
| | | 32 | 2.5e-4 | 0.1 | 78.88 ACC |
| | ASA | 16 | 2.5e-4 | 0.0 | 66.09 PCC |
| | | 16 | 2.5e-4 | 0.05 | **66.37** PCC |
| | | 16 | 5.0e-4 | 0.05 | 65.98 PCC |
| | | 16 | 2.5e-4 | 0.1 | 66.23 PCC |
| | | 32 | 2.5e-4 | 0.1 | 66.31 PCC |
| | PPI | 8 | 1.25e-4 | 0.0 | 61.82 AUC ROC |
| | | 8 | 2.5e-4 | 0.0 | 57.91 AUC ROC |
| | | 8 | 1.25e-4 | 0.05 | **62.11** AUC ROC |
| | | 32 | 2.5e-4 | 0.05 | 58.68 AUC ROC |
| | | 8 | 1.25e-4 | 0.1 | 60.72 AUC ROC |
| | EPI | 8 | 6.25e-5 | 0.05 | 68.31 AUC ROC |
| | | 16 | 2.5e-4 | 0.05 | **69.90** AUC ROC |
| | | 16 | 5.0e-4 | 0.05 | 68.47 AUC ROC |
| | | 8 | 1.25e-4 | 0.1 | 68.68 AUC ROC |
| | HPR | 16 | 2.5e-4 | 0.0 | 18.14 PCC |
| | | 32 | 6.25e-5 | 0.0 | **19.61** PCC |
| | | 8 | 2.5e-4 | 0.1 | 18.81 PCC |
| | | 32 | 6.25e-5 | 0.1 | 19.18 PCC |
| `base` | SS3 | 8 | 2.5e-4 | 0.0 | **74.84** ACC |
| | | 8 | 5.0e-4 | 0.0 | 73.48 ACC |
| | | 8 | 2.5e-4 | 0.05 | 74.77 ACC |
| | | 8 | 5.0e-4 | 0.05 | 66.25 ACC |
| | | 8 | 5.0e-4 | 0.1 | 74.83 ACC |
| | SS8 | 8 | 2.5e-4 | 0.0 | **61.89** ACC |
| | | 8 | 5.0e-4 | 0.0 | 53.52 ACC |
| | | 8 | 2.5e-4 | 0.05 | 61.70 ACC |
| | | 4 | 2.5e-4 | 0.0 | 60.70 ACC |
| | BUR | 4 | 2.5e-4 | 0.0 | 79.22 ACC |
| | | 8 | 2.5e-4 | 0.0 | 79.49 ACC |
| | | 8 | 2.5e-4 | 0.05 | 79.38 ACC |
| | | 8 | 2.5e-4 | 0.1 | **79.55** ACC |
| | ASA | 8 | 1.25e-4 | 0.0 | **66.07** PCC |
| | | 8 | 6.25e-5 | 0.05 | 64.98 PCC |
| | | 8 | 1.25e-4 | 0.05 | 65.73 PCC |
| | | 4 | 1.25e-4 | 0.1 | 65.10 PCC |
| | | 8 | 1.25e-4 | 0.1 | 65.89 PCC |
| | | 8 | 1.25e-4 | 0.2 | 65.10 PCC |
| | PPI | 8 | 6.25e-5 | 0.0 | **61.62** AUC ROC |
| | | 8 | 1.25e-4 | 0.0 | 60.22 AUC ROC |
| | | 4 | 6.25e-5 | 0.05 | 60.78 AUC ROC |
| | | 4 | 1.25e-4 | 0.1 | 60.49 AUC ROC |
| | | 8 | 6.25e-5 | 0.1 | 59.29 AUC ROC |
| | | 4 | 1.25e-4 | 0.2 | 60.67 AUC ROC |
| | EPI | 8 | 6.25e-5 | 0.0 | 69.47 AUC ROC |
| | | 4 | 6.25e-5 | 0.05 | 69.27 AUC ROC |
| | | 8 | 6.25e-5 | 0.1 | 69.81 AUC ROC |
| | | 8 | 1.25e-4 | 0.1 | **69.51** AUC ROC |
| | HPR | 8 | 6.25e-5 | 0.0 | 18.67 PCC |
| | | 8 | 6.25e-5 | 0.1 | **18.72** PCC |
| | | 8 | 1.25e-4 | 0.1 | 18.55 PCC |
| | | 4 | 6.25e-5 | 0.2 | 18.58 PCC |

The hyperparameters are tuned on each structural prediction task individualy on the validation set. First, all hyperparemeter combinations were tested once in a grid search. For each tasks, the four till eight best combinations were further investigated by training and validate the structural prediction tasks another three times. The mean performance for the selected hyperparameter combinations per tasks are shown here. The highest performance is shown in bold.

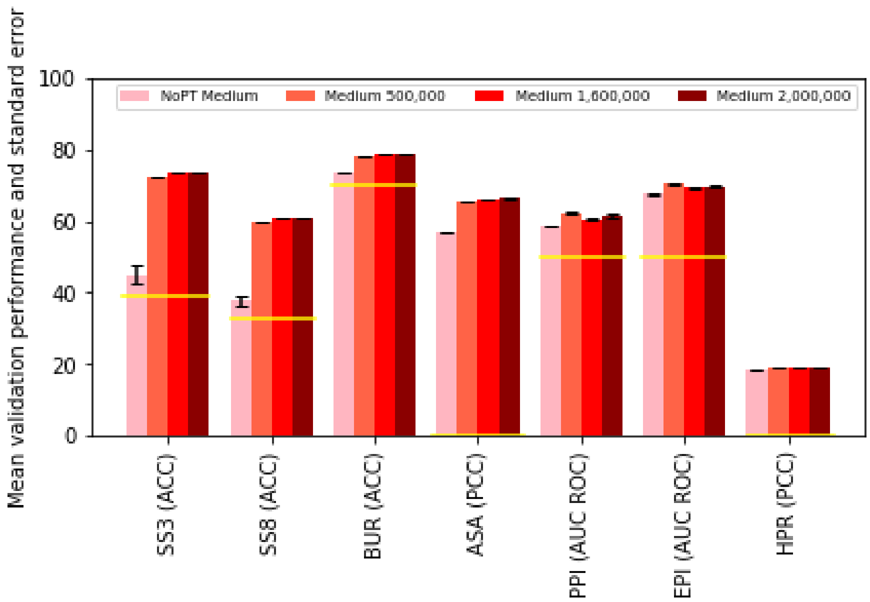


**A**

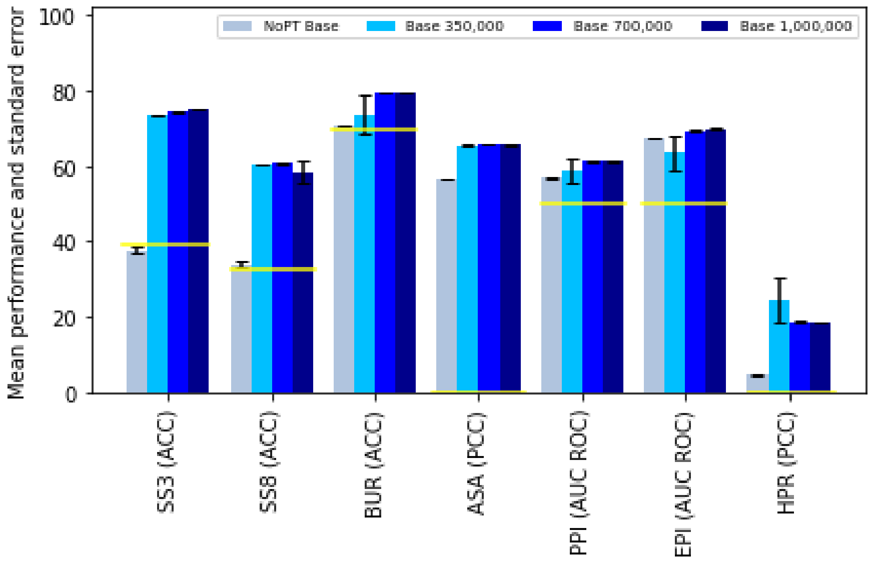


**B**

**Figure 2.** *Performance of fine-tuning tasks monitored during pre-training.* Prediction performances of the validation set, (A) for a medium sized model, and (B) for a base sized model, trained without the pre-training step (pink/grey), and on a pre-trained base size model for 350 000 (light red/blue) steps, 700 000 steps (red/blue) and 1 000 000 steps (dark red/blue).
